# A systematic review of social contact surveys to inform transmission models of close contact infections

**DOI:** 10.1101/292235

**Authors:** Thang Van Hoang, Pietro Coletti, Alessia Melegaro, Jacco Wallinga, Carlos Grijalva, W. John Edmunds, Philippe Beutels, Niel Hens

## Abstract

Social contact data are increasingly being used to inform models for infectious disease spread with the aim of guiding effective policies on disease prevention and control. In this paper, we undertake a systematic review of the study design, statistical analyses and outcomes of the many social contact surveys that have been published. Our primary focus is to identify the designs that have worked best and the most important determinants and to highlight the most robust findings.

Two publicly accessible online databases were systematically searched for articles regarding social contact surveys. PRISMA guidelines were followed as closely as possible. In total, 64 social contact surveys were identified. These surveys were conducted in 24 countries, and more than 80% of the surveys were conducted in high-income countries. Study settings included general population (58%), schools/universities (37%) and health care/conference/research institutes (5%). The majority of studies did not focus on a specific age group (38%), whereas others focused on adults (32%) or children (19%). Retrospective and prospective designs were used mostly (45% and 41% of the surveys, respectively) with 6% using both for comparison purposes. The definition of a contact varied among surveys, e.g. a non-physical contact may require conversation, close proximity or both. Age, time schedule (e.g., weekday/weekend) and household size were identified as relevant determinants for contact pattern across a large number of studies. The surveys present a wide range of study designs. Throughout, we found that the overall contact patterns were remarkably robust for the study details. By considering the most common approach in each aspect of design (e.g., sampling schemes, data collection, definition of contact), we could identify a common practice approach that can be used to facilitate comparison between studies and for benchmarking future studies.

## Introduction

Despite the great progress in infectious disease control and prevention that were initiated during the last century, infectious pathogens continue to pose a threat to humanity, as illustrated by SARS, influenza, antimicrobial resistant bacteria, Ebola, and resurgent measles, disrupting everyday life, burdening public health and dominating media headlines well into the 21st century.

Many infectious diseases can spread rapidly between people within and between age groups, households, schools, workplaces, cities, regions and countries through a diversity of social contacts^1^. Understanding and quantifying social mixing patterns is therefore of critical importance to establish appropriate simulation models of the spread of infectious diseases. Such mathematical transmission models have become indispensable to guide health policy. Which interventions should be offered to which people in which circumstances? How would such interventions affect transmission chains and the disease burden throughout the population? What would be the population effectiveness and cost-effectiveness of such interventions? Well-informed answers to these questions require mathematical models. The validity of such models depends heavily on the appropriateness of their structure and their parameters, including what they assume about how people interact.

Indeed, a transmission model’s integrated mixing patterns (i.e., who mixes with whom?) have a strong influence on the transmission parameters (i.e., who infects whom?). The latter are the most influential drivers for the outputs of such models. Whereas 20th century models made strong assumptions about mixing patterns, it has become increasingly common to use empirical data on social interactions as a direct model input over the last decade. For sexually transmitted infections, data from surveys on sexual behavior were available for use. For infectious diseases that are transmitted by direct contact, minimal data on relevant social contacts was available for use. A study that aimed to collect precisely this information was conducted using a convenience sample^2^. This study was followed by a study that reported on relevant social contacts in a representative sample of the population that covered all ages in a town^3^. The landmark study that reported on relevant social contacts in representative samples for eight different European countries using contact diaries was the POLYMOD study^4^. Numerous other studies have been reported since. Several of these studies report on social mixing patterns as obtained through direct observation, contact diaries or electronic proximity sensors. The strengths and weaknesses of these methods have been discussed^5^. Nevertheless, a comprehensive review of the study designs for contact diaries, statistical analyses of social contact data and major determinants of mixing patterns is lacking for this rapidly growing field of research, a gap which we aim to fill here.

In the current paper, we systematically retrieve and review the literature on social contact surveys. First, we provide an overview of the literature to help identify a standard. Second, we present the different approaches for data collection and identify strength and limitations. Third, we report on the main determinants of contact. We use these findings to guide future studies.

## Materials and methods

We conducted a systematic review in accordance with the Preferred Reporting Items for Systematic Reviews and Meta-Analyses (PRISMA) guidelines^6^.

### Search strategy

We queried PubMed and ISI Web of Science, without time and language restriction up to Jan 31, 2018 using the following search string:

*(([survey*] OR [questionnaire*] OR [diary] OR [diaries]) AND ([social contact*] OR [mixing behavio*] OR [mixing pattern*] OR [contact pattern*] OR [contact network*] OR [contact survey*] OR [contact data])*

EndNote X7 was used to eliminate duplicates and manage the search results^7^.

### Inclusion criteria

Studies were considered eligible if they fulfilled all of the following criteria: (1) primary focus on face-to-face contacts of humans, implying the physical presence of at least two persons during contact; (2) contacts relevant for the transmission of close-contact infections; (3) contacts recorded using a diary-type system on paper or in electronic format; (4) full-text version available.

### Exclusion criteria

Studies were excluded if they involved at least one of the following: (1) primary focus on human-animal or animal-animal contacts; (2) recording contacts exclusively relevant for sexually transmitted, food-, vector- or water-borne diseases; (3) using exclusively proximity sensor devices or observational methods to collect contact data; (4) including contacts without physical presence (e.g., phone, internet/social media contacts) or without the possibility to distinguish them; (5) recording the frequency or regularity but not the number of contacts over a given time period; (6) meeting abstracts, books, theses or unpublished journal articles.

An overview of the selection process is presented in Fig 1. Title, abstract and full text were screened initially by the first author and double-checked by the second author in case of doubt.

**Figure 1:**
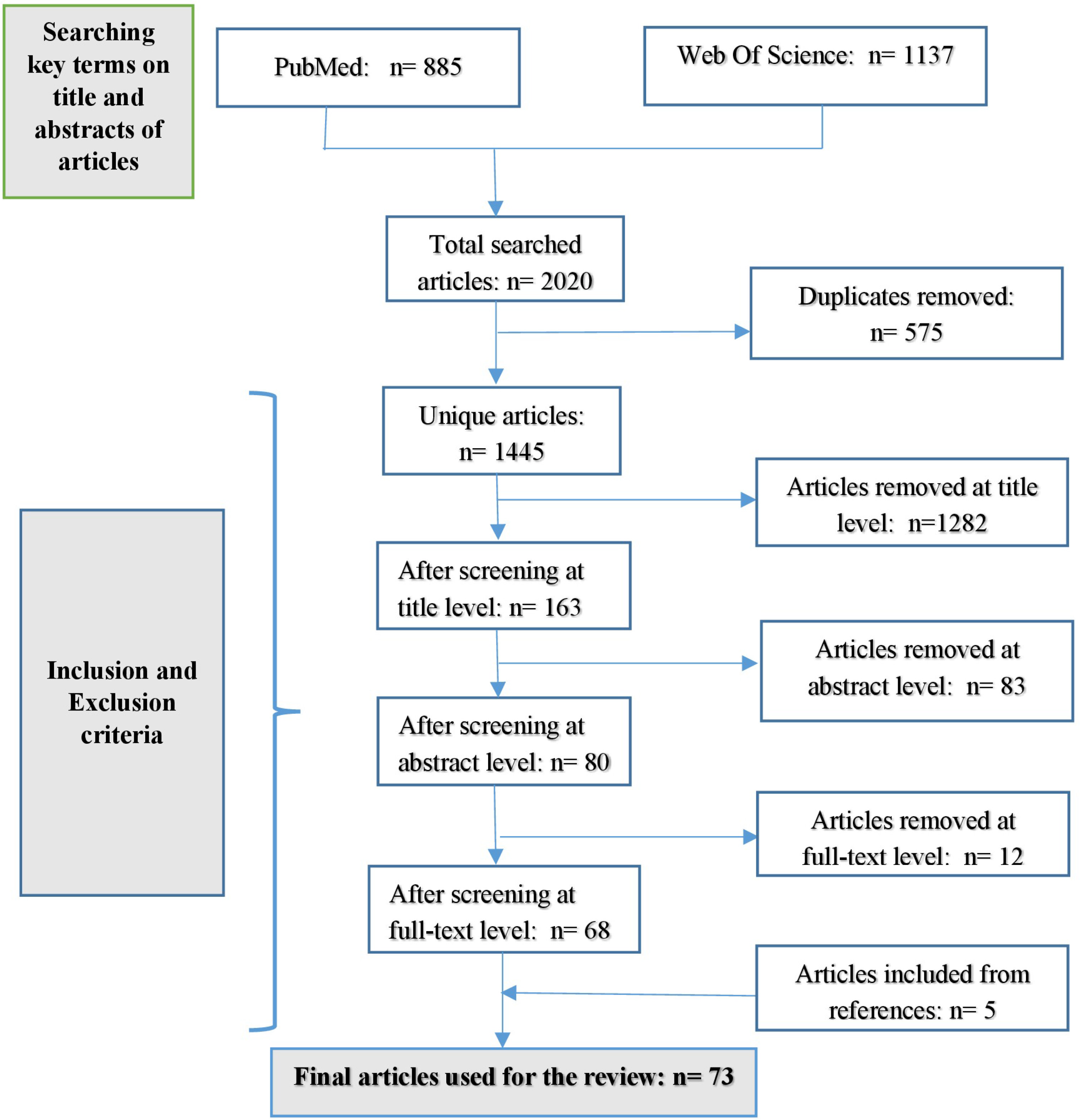
The PRISMA flowchart of the search process.

## Results

### The screening process

After removing 575 duplicates, a total of 1445 unique articles were identified. Using the inclusion and exclusion criteria, 1282 articles were removed at title level and an additional 83 articles were removed at abstract level, yielding 80 articles for full-text screening. At this stage, an additional 12 articles were excluded: 9 did not quantify the number of contacts made in a specific time period, 2 used contact definitions that did not require physical presence and 1 focused exclusively on proximity sensor devices. Using the reference lists of the 68 remaining articles, we identified 5 additional articles. Thus, a total of 73 articles were identified for review (see Fig 1 for more details).

### Country settings

The 73 remaining articles covered 64 social contact surveys conducted in 24 countries spread over 5 continents: 12 European (Belgium^4,8–10^, Finland^4^, France^4,11,12^, Germany^4,13–16^, The United Kingdom (UK)^2,4,17–25^, Italy^4^, Luxembourg^4^, Poland^4^, Sweden^26^, Switzerland^27,28^, The Netherlands^3,4,9,29,30^, and The Russian Federation^31^), 5 Asian (China^32–34^, Japan^35^, Taiwan^36^, Thailand^29^ and Vietnam^37^), 4 African (Kenya^38^, South Africa^39,40^, Zambia^39^ and Zimbabwe^41^, 2 American (Peru^42^ and The United States of America^43–47^) and 1 Oceanian country (Australia^48–50^). More details on number of social contact surveys in each of these countries are shown in the global map (see S7 Fig). Only 14 studies were conducted at the whole-country level ^4,11,12,24,26,30,35,36^, whereas remaining studies focused on a region^10,32,33,37,39,41,45^, a city/town^3,34,38,40,49^ or a specific setting (school/university, health care facility, etc.) and were therefore not representative of the entire country. Fig 2 demonstrates that 40 out of 64 the surveys were conducted in Europe followed by Asia with 10 surveys. In contrast, only a few surveys were conducted in other regions. In this representation, we count several countries separately, even if they were included as part of a single project^4,9,29,39^.

The number of surveys greatly increased over time from only 4 surveys before the year 2000 up to 37 surveys after 2009, indicating that social contact surveys are increasingly conducted. In addition, no survey was conducted outside Europe before 2005. One survey did not indicate the year it was conducted. For this study we used the publication year minus two as a proxy^2^.

### Study settings and subjects

Greater than half of the social contact surveys were conducted in the community/general population (58%). Of these, there were only 4 household-based surveys that asked every member of each participating household to complete the survey^12,33,37,42^, whereas in other surveys only one person in each participating household was asked to participate. The majority of surveys conducted in the general population aimed at people of all ages (65%) ^4,10,12,24–26,30–32,34–38,40–42,45^. In contrast, two surveys excluded infants younger than one year^3^, and one excluded children less than two years^33^. Four surveys focused exclusively on adults^27,39,39,49^, two survey investigated contact patterns of infants (under 11 weeks 23 and under 1 year old^50^), and one survey aimed at patients with pandemic influenza AH1N1 (swine flu)^18^. More specific settings of schools or universities constituted 38% of the surveys, of which 11 surveys were conducted at schools (primary schools^14^, secondary schools^21^, high schools ^46,47,51–54^ and together^18,19,44^) and 13 surveys were performed in universities ^2,8,15,17,22,29,31^. Of surveys performed in schools/universities settings, in addition to students serving as main study subjects, staff, family and friends of students were also asked to participate in 6 surveys^2,8,9,22,47^. In addition to school/university settings, we also identified 1 contact survey on nurses in a health-care setting^13^, 1 survey at a conference^16^ and 1 survey in a research institute ^28^. In addition, there were 4 web-based surveys open for anyone to participate, and respondents of these surveys, namely Internet users, are considered as a separate group of study subjects ^20,24,55^.

### Sample size and response rate

Among social contact surveys conducted in the general population, the smallest survey only consisted of 54 participants in Switzerland^27^, and the largest survey consisted of 5388 participants in the UK^24^. The largest survey in a school/university setting contained 803 participants in Germany^15^ (see S1 Fig). The response rate was reported in 36 out of 64 surveys and ranged from 4% in population based surveys^24^ up to 100% in a school-based survey^15^. Of these 34 surveys, only 3 considered the response rate beforehand to estimate the sample size^13,36,45^. Fu et al.^36^ referred to the average response rates from various nationwide surveys over the past 5 year. These researchers set a goal of 1900 but used a sample of 4207 based on a response rate of 45% for general population-based surveys. DeStefano et al.^45^ determined survey response rates using the Council of Survey Research Organization (CASRO) method, leading to 4135 completed interviews, exceeding the goal of 4000. Bernard et al.^13^ also assumed a dropout rate of 20% of the initial participants and accounted for that proportion of declining participants. Instead of considering the response rate, some surveys established criteria to substitute those who refused or were not reached after several attempts^37–39^. For example, in a survey conducted by Horby et al.^37^, if a selected household declined to participate, the nearest neighbor was approached to participate in the survey. Only one survey^24^ presented details on biases in the demographic characteristics of respondents, such as age, gender and household composition. Specifically, males between 60 and 90 years of age are more likely to appear in the sample, compared with the population distribution; one- and two-person households are overrepresented. Twenty surveys determined sampling weights based on demographic characteristics of the populations to reduce the effects of sample bias ^4,9–11,23,24,30,34–39,49^.

**Figure 2:**
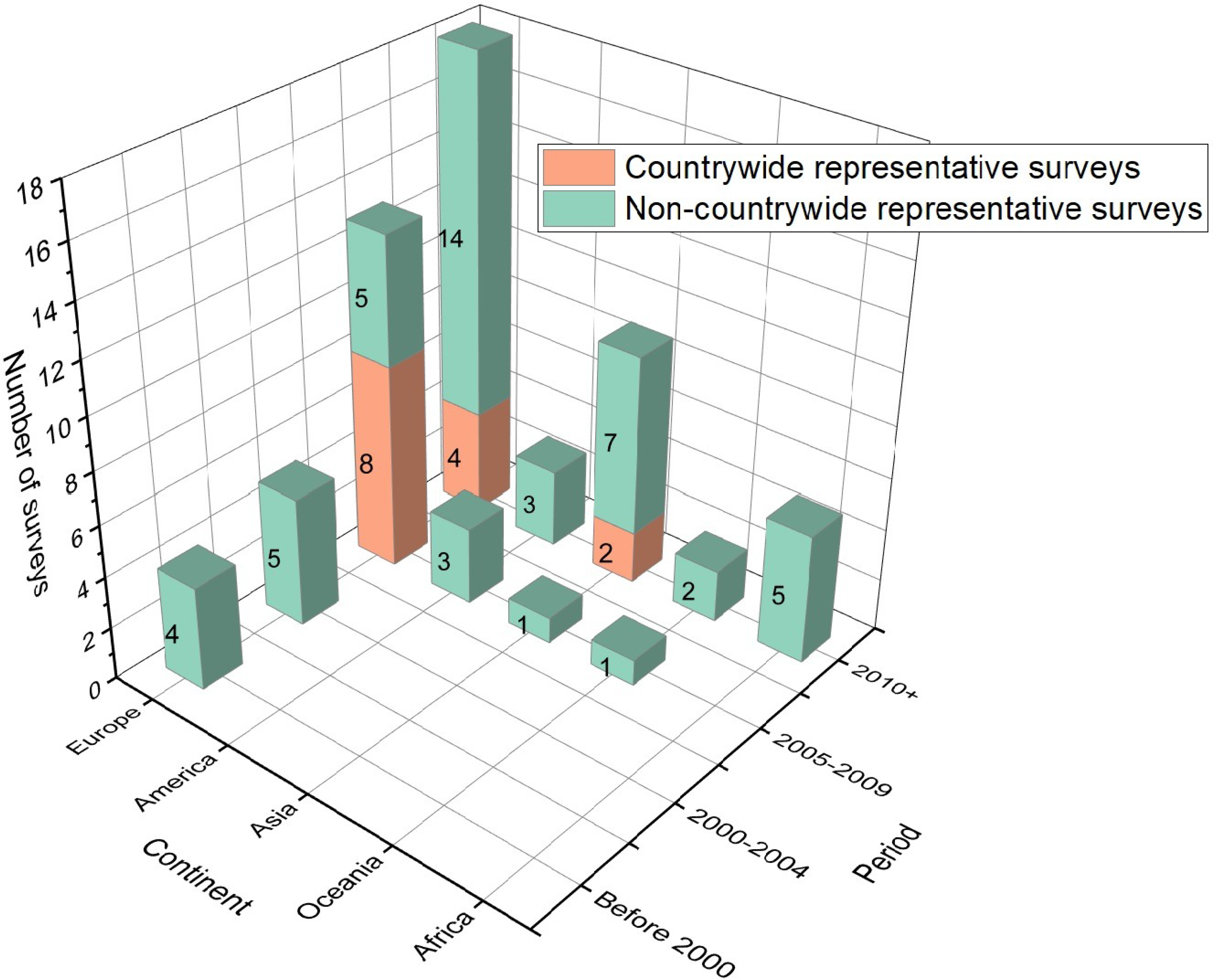
Distribution of number of social contact surveys. Distribution of number of representative surveys based on continents and time periods.

### Sampling methods

Approximately half (44%) of the surveys employed convenience sampling, in which subjects were selected based on of their convenient accessibility and proximity to researchers^56^. This sampling technique was also used for the sake of comparing data collection tools^16,47,48,54,57^, data collection methods^58^ or study designs^8,15,48^.

Other sampling techniques can generate samples that can be considered representative for a population but to different extents. Given that random sampling is inherently difficult to implement and generally expensive^56^, this technique was used in 7 surveys^3,24,26,32,33,49^. Among these studies, only 2 surveys were considered representative of the entire country^24,26^, and the remaining surveys were representative of a region^32,33^ or a city/town^3,49^. Given that it is easier to implement and remain representative, multi-stage and stratified sampling were employed in 3 surveys^13,36,37^ and 10 surveys^10,12,23,35,38–40,45^, respectively. In addition, 10 surveys relied on quota sampling, which aimed to represent certain characteristics of a population, e.g., age, sex, geography, etc. Of these surveys, 9 were conducted at the whole-country level^4,11,30^, and one survey focused on one specific region^34^. In addition, one survey used mixed samplings, in which a convenient sample of students was obtained in 2 schools and a random sample of the general population was obtained in one province^31^. When restricted to surveys conducted in the general population, the sampling frames were obtained from a population register/post office address database ^4,26,30,33,36–38,40,42^, landlines and mobile numbers^4,10–12,45,49^ or a list of participants of a larger study^32,35,39,41^. Five surveys used an online respondent-driven method, which can be considered as a snowball/chain sampling technique^29,43,55^. In particular, these surveys used online systems to recruit “seed” participants who were asked to recruit people they know into the surveys. This process was performed until the desired sample size was obtained. Only one survey did not state information on sampling techniques^32^. Finally, 3 surveys conducted at the general population level used a convenience sample^27,42,48^, therefore not relying on a sampling frame. More details on the distribution of sampling schemes based on time and regions are presented in S2 Fig.

### Study Design

By prospective design, we mean that respondents are informed in advance of the day(s) that they are requested to record their contacts^4,15,30^. In a retrospective design, respondents recall their contacts over a past time period without prior warning or instruction that they would be requested to do so. Of 64 surveys, 29 (45%) used a retrospective design, and 26 (41%) used a prospective design. Only 4 surveys (6%) used both designs for the purpose of comparison^8,15,48^. For five surveys (8%), it was not completely clear whether the study was prospective or retrospective^18,22,28,46^. S4 Fig displays the trend of using study designs in social contact surveys over time, revealing that the retrospective design was more favored by researchers, except in the period 2005-2009 in which eight prospective surveys were implemented under the same funding source from the European commission project POLYMOD^4,30^. Of the 14 country-wide surveys, the prospective design was employed in 10 surveys,^4,11,24,30^ and a retrospective design was employed in 4 surveys^12,26,35,36^.

Mccaw et al.^48^ compared the prospective design with the retrospective design in the same survey population with a small convenience sample of adult participants. These researchers asked respondents to report their social contacts in 3 typical defined days and compared these data with the results in the same days in the following week, revealing that the prospective design yielded more complete data and more contacts (mean 27.5 versus 15.0). The difference in number of recorded contacts between these two study designs was less pronounced in the work of Mikolajczyk et al. conducted among school children^15^. On average, students recorded two more contacts in the prospective part compared with the retrospective part (asking about a day preceding the survey day) for workdays and one more contact for weekends. Beutels et al^8^ observed statistically non-significant differences in average number of contacts made when respondents were asked prospectively on a random day or retrospectively about “yesterday”. In this study, 64% of respondents indicated however that the retrospective design was difficult or very difficult, whereas 65% found the prospective diary-based approach to require minimal effort.

### Data collection tools

Most surveys (83%) used paper diaries to collect contact data. Diaries were delivered and collected by mail in 9 surveys^4,10,11,18,24^ and in person in the remaining surveys. Six surveys exclusively used online diaries^9,20,29,43^, and 4 surveys used both online and paper diaries^8,24,34,35^. A Personal Digital Assistant (PDA) was used with paper diaries in one survey in 2008^48^. In contrast, proximity sensor devices were used with online diaries in 1 survey in 2012^47^ and with paper diaries in 2 surveys in 2013^54^ and 201^16^. Most of these studies primarily aimed to explore different social contact data collecting tools.

In contrast to the work of Beutel et al.(2006)^8^, which concluded that the paper diary yielded similar results of contact numbers compared with the online diary, Leung et al.^34^ showed that participants using paper diary reported on average 9.99 contacts per day, which is substantially increased compared with those using the online diary with only 5.10 contacts per day. In McCaw et al.^48^, respondents preferred the paper diary compared with the PDA based on timeliness and completeness. The majority of respondents (63%) described the paper diary as “easy” to use, whereas only 35% respondents had the same opinion regarding the use of PDA. Smiesziek et al.^47^ conducted a survey at a high school using both the online diary and wearable proximity sensors, making the two sources of data comparable by matching names of contacts reported by online diaries with names associated with sensor ID numbers. They found the use of online diaries to be more accurate for longer duration contacts but much less accurate for short duration contacts. Mastrandrea et al.^54^ and Smiesziek et al.^16^ reached similar conclusions comparing paper diaries with wearable proximity sensor devices, with better accuracy for contacts of longer duration using paper diaries. However, there was a distinction in self-reported ease of use with 25% of respondents reporting difficulties in remembering contacts to complete paper diaries, and 25% stating that filling in the diary was too much work. In contrast, 93% respondents felt comfortable having their contacts measured by sensors^16^.

### Data Collection methods

Fifty-two of the 64 surveys (81%), relied on self-reporting, i.e. respondents single-handedly completed a paper or online diary after receiving oral instructions (in person or via telephone) or written guidelines from investigators. A face-to-face interview was used in nine surveys ^3,12,32,33,36,37,39,42^ and a Computer-Assisted Telephone Interview (CATI) was employed in three^26,45,49^. Although self-reporting can be used for both retrospective and prospective designs, face-to-face interviews and the CATI only allow for a retrospective design, unless they are supplemented by some form of personal diary keeping that is used during the interview. Akakzia et al.^58^ found the paper diary to provide more number of contacts compared with CATI (the median number of reported daily contacts per participant was 13.5 for the paper diaries and 4 for CATI) because it gives the respondent more time to recall contacts. All 4 household-based surveys conducted face-to-face interviews at the respondent’s residence^12,33,37,42^. Face-to-face interviews exhibited a greater response rate (46%^36^ to 100%^42^) compared with self-reporting and CATI. We could not identify any contact survey directly comparing self-reporting with a face-to-face interview, but different surveys were compared in one work^3^ .

### Definition of contact

Forty (63%) surveys distinguished physical and non-physical contacts. Physical contacts were consistently defined as involving any sort of skin-to-skin touching, e.g., handshake, hug, kiss, etc. The definition of non-physical contacts differed somewhat among surveys. Specifically, the majority of surveys using two types of contacts defined a non-physical contact as a two-way conversation of at least 3 words at a distance that does not require raising one’s voice ^4,8,34,37,41,48,50,52,58^. In some other surveys, the definition involved close proximity (e.g. verbal communication made within 2 meters) without specification of a minimum number of words to be exchanged^11,24,28,36,39,59^. Of note, that since the POLYMOD contact studies were executed^4^, its contact definition was applied in several subsequent surveys^31,34,37,41,42,50,53^, both for physical and non-physical contacts. Fifteen surveys used only one type of contact with varying definitions either involving a face-to-face conversation^2,3,14,21,31,43^ or being in close proximity within a certain distance (e.g., within one arms length)^29,46,54^, both regardless of any skin-to-skin touching^35,47^ or only involving direct skin-to-skin touch^26,38^. Nine surveys used more than two types of contact^15,17,27,44^. Only one survey attempted to record casual contacts occurring in an indoor location without the requirement for a conversation or any type of touch^40^. Eight remaining surveys added kissing or intimate sexual contact as different types of contacts^15,17,27,44^ or asked respondents to record contacts made in small/large groups or occasional contacts within 2 meters in local transportation or crowded places separately^15^.

### Reporting time period

Greater than half the surveys asked respondents to report contacts they made during a single day, whereas only six surveys used a reporting time period of greater than 3 days. A reporting day was defined in most of these surveys as being either (1) between 5 A.M and 5 A.M the next morning or (2) the time period between getting up and going to bed. The longest time period identified is 3 days in a prospective survey^17,48^ and 10 weeks in a retrospective survey^43^. Some surveys recorded both weekdays and weekend days^4,8,17,34,51–53^. These researchers observed marked day-of-week variation. An increased frequency of contacts occur during weekdays with respect to weekends is generally observed. In contrast, the Hong Kong study^34^ identified an age-dependent behavior with children and adolescents under 18 also reporting more school contacts during weekends than weekdays. This peculiar behavior is linked to the specific Hong Kong schooling system, in which students participate in many extra-curricular activities with their schoolmates during weekend^34^. Similarly, several surveys, which were conducted on school-based populations, explored differences in contact patterns between school term or regular periods versus holiday periods^4,18–21,41^. The results from these surveys indicated that school closure is associated with large reductions in the number of contacts and would therefore have a substantial impact on the spread of a newly emerging infectious disease^21,60^. Eames et al^18^ quantified the changes in social contact patterns experienced by individuals experiencing an episode of Influenza A(H1N1) on two randomly assigned days: one day while being ill and one day when recovered.

### Characteristics of participants and contactees

Most surveys collected a range of demographic background characteristics of study participants, e.g., age, gender, education and household size. Some surveys also asked participants to record any influenza-like-illness symptoms they experienced on the day of surveying^29,51,55^ or whether their day was in any way special (due to holiday, sickness, etc.)^4,10^.

The characteristics of contactees required greater effort given that participants had to remember and/or record anyone they met, even if they were vague acquaintances or first time fleeting encounters. Among the characteristics of contactees, age and gender are considered to be the most important determinants of the mixing patterns given that they can help explain age and gender differences in the epidemiology of infectious diseases^11^. In some surveys, if the participants did not know the exact age of their contacts, they were asked to provide or choose an age range (the mid-point of this range was often used for the purpose of analysis)^2,4,11,30,31,37,53^. Thirty six of 64 (56%) surveys recorded both age and gender of contactees, and 16(25%) surveys recorded only age of contactees. In contrast, 7 (11%) surveys required participants to simply report the number of different contactees without recording any of their characteristics^15,27,43,45,47^. In 5 surveys (8%), it was not clear what contactee characteristics participants had to report^22,24,28,33,46^. Along with age and gender, several surveys also asked participants to record health status of contactees and any symptoms they experienced, e.g., coughing, sneezing, fever, etc.^43,51–53^ or whether they wore a protective mask^51,52^. Most of these surveys were conducted in school or university settings using a convenience sample.

### Information about contacts

Along with information on types and intimacy of contacts (physical or non-physical contacts), the location, duration and frequency of contacts were also studied. Participants were asked to record information about location, duration and frequency of each contact in 77%, 67% and 52% of contact surveys, respectively. If the survey participant met the same person more than once in different contexts during the day, then they were asked to estimate the summed duration and report all of the locations^2,4^ . Thirty-four surveys recorded both location and duration. In addition, 29 surveys recorded both location and frequency, and 31 surveys recorded both duration and frequency. All these contact characteristics were jointly recorded in 27 surveys (42%). In addition to this information, several surveys also asked participants to report distance from home of contacts, which could help provide more insight into how infectious diseases spread regionally^10,23,24,33,36^.

### Mean number of contacts and analysis of determinants

**Figure 3:**
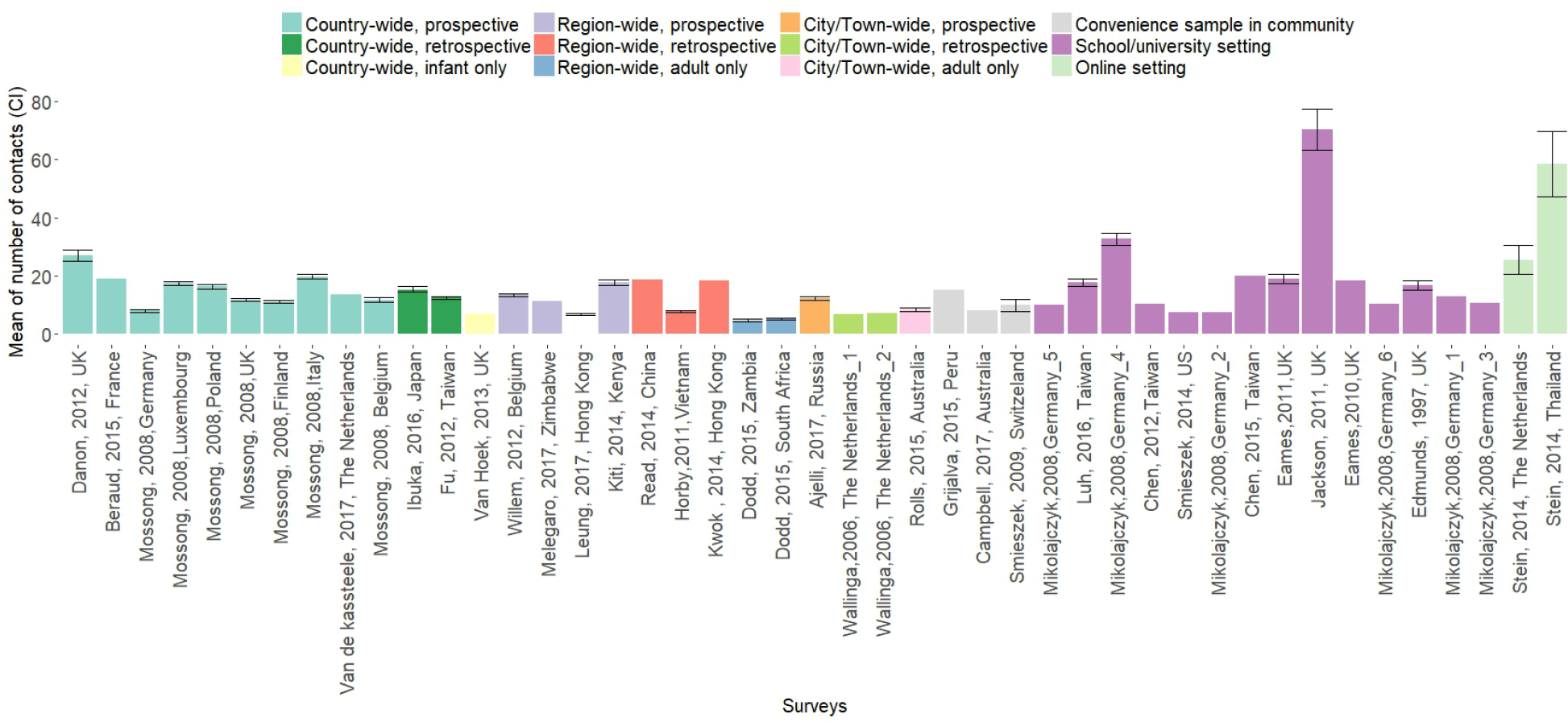
Average number of contacts among social contact surveys. Average number of contacts measured and 95% CI. For surveys reporting mean number and standard deviation a 95% CI for the mean was computed. Surveys are labeled according to the publication’s first author, year and to the country in which the survey was performed. Ordering is performed based on increasing sample size within the specific design strata.

Of the 64 surveys, 45 explicitly reported the average number of contacts measured without any stratification (Fig 3). To compare these survey results, we categorized them into 12 groups with different extents of representativeness (for country, region or town/city), study designs and settings. In country-wide prospective surveys, the average number of contacts ranges from a minimum of 7.95 ([7.61, 8.29] 95% CI) in Germany^4^ to a maximum of 26.97 ([25.05, 28.89] 95% CI) in the UK^24^. In country-wide retrospective surveys, these values ranges from 12.5 ([12.09, 12.91] 95% CI) in Taiwan^36^ to 15.3 ([14.4, 16.3] 95% CI) in Japan^35^. Six surveys conducted in the general population asked participants not to report details of professional contacts in the diary but to provide an estimate and age distribution if they had more than 10 contacts (surveys in Finland^4^, Germany^4^ and the Netherlands^4^) or more than 20 professional contacts (surveys in Belgium^4,10^ and France^11^). The additional professional contacts are not included in calculation of means presented in Fig 3. Among school/university-based surveys, the highest number of contact (70.3 [63.23,77.37] 95%CI) was observed in a secondary school in the UK^21^. Only sixteen surveys reported median and quantiles of contacts without any stratification (see S6 Fig).

Fig 4 presents the number of surveys that have analyzed possible determinants for the number of contacts. For every determinant, we report the number of surveys that identified a relevant connection with the number of contacts, the number of surveys that did not identify such a connection and the number of surveys that did not delve into the matter. Strong evidence is identified for the age (34 yes vs. 5 no) and the household size (21 yes vs. 4 no) of the participant to affect the number of contacts. Only 5 surveys identified gender as a relevant indicator for the number of social contact, in contrast to 23 surveys that did not identify a significant relation. Social contacts are also affected from the daily routine (29 yes vs. 6 no) with a larger number of contacts during weekdays compared with the weekend (S5 Fig a, with the exception of^34^). Similar results hold for term time versus holidays with all of the 8 surveys analyzing the issue identifying a larger number of contacts during term time (S5 Fig b). In addition, a self-reported healthy status is associated (5 yes vs. 1 no) with a larger number of contacts with respect to feeling ill (S5 Fig c). For example, Eames et al.^18^ found that respondents made approximately two-thirds fewer contacts when they were not feeling well compared with when they were feeling well. The relation between social contacts and urbanization has been analyzed in 3 surveys. One survey found a larger number of contacts in peri-urban areas compared with rural areas^41^, one found the opposite^38^, and one did not find any evidence^33^. Finally, two surveys analyzed contacts during and outside flu seasons. Of these surveys, one survey^45^ used a model to adjust for other factors, e.g., age and sex, and one did not^53^. However, both surveys identified no relevant effect (S5 Fig d). Table 1 provides summaries of all 64 social contact surveys.

**Figure 4:**
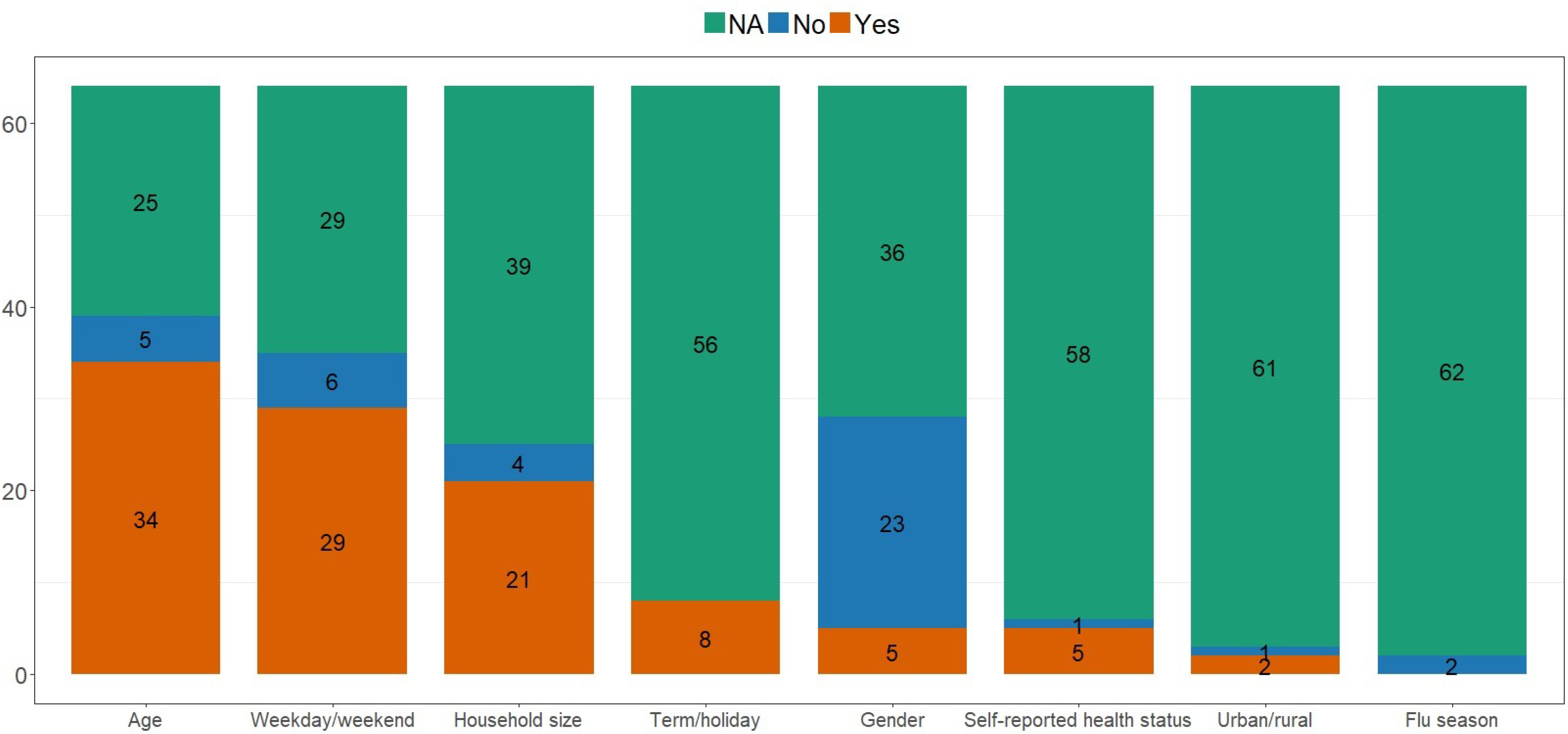
Determinants of number of social contacts. Determinants of number of social contacts. Surveys are tagged as ‘Yes’ if a relevant connection between the number of contacts and the determinant was identified, ‘No’ if evidence was not identified, or ‘NA’ if the given determinant was not analyzed.

**Table 1:**
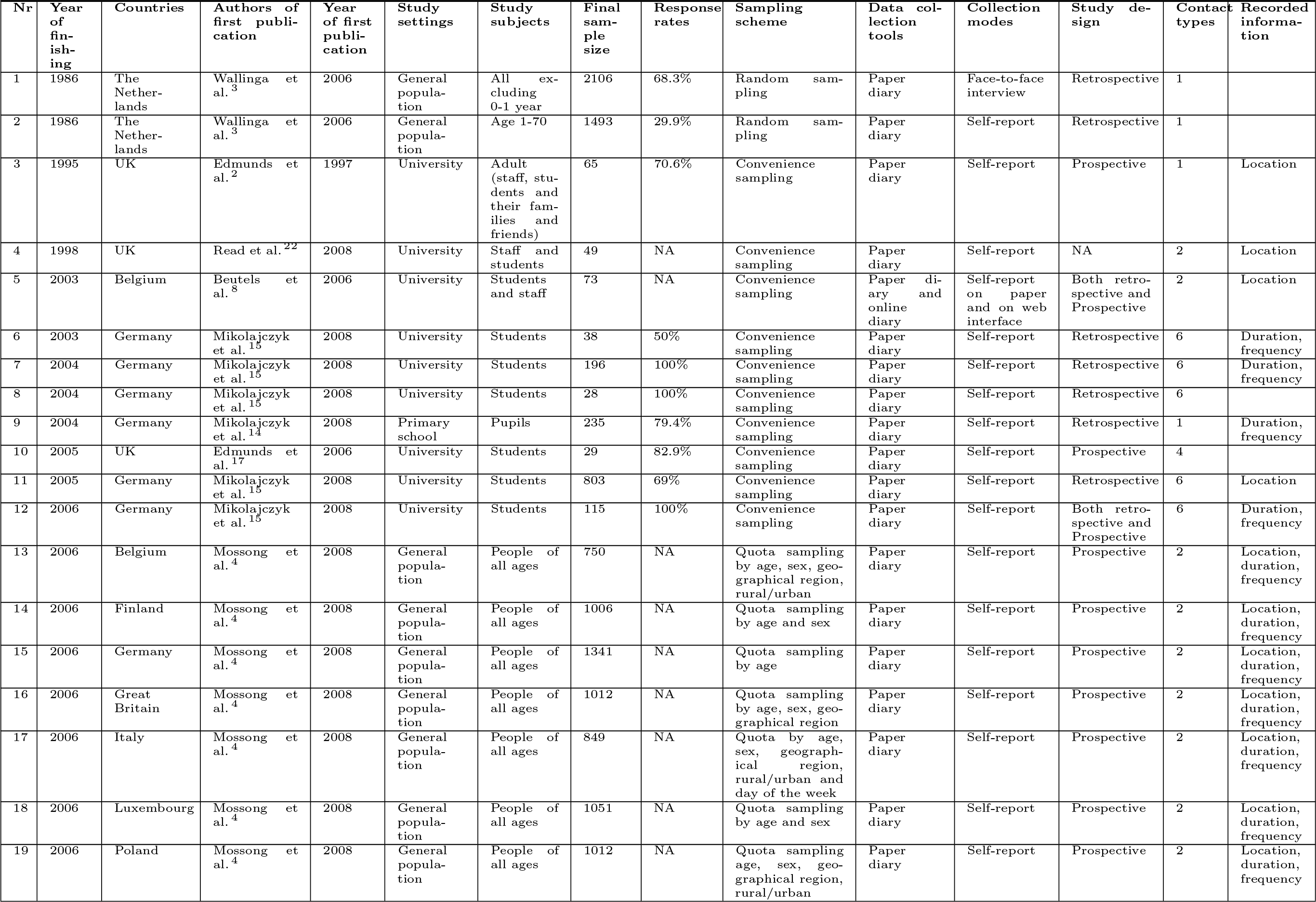
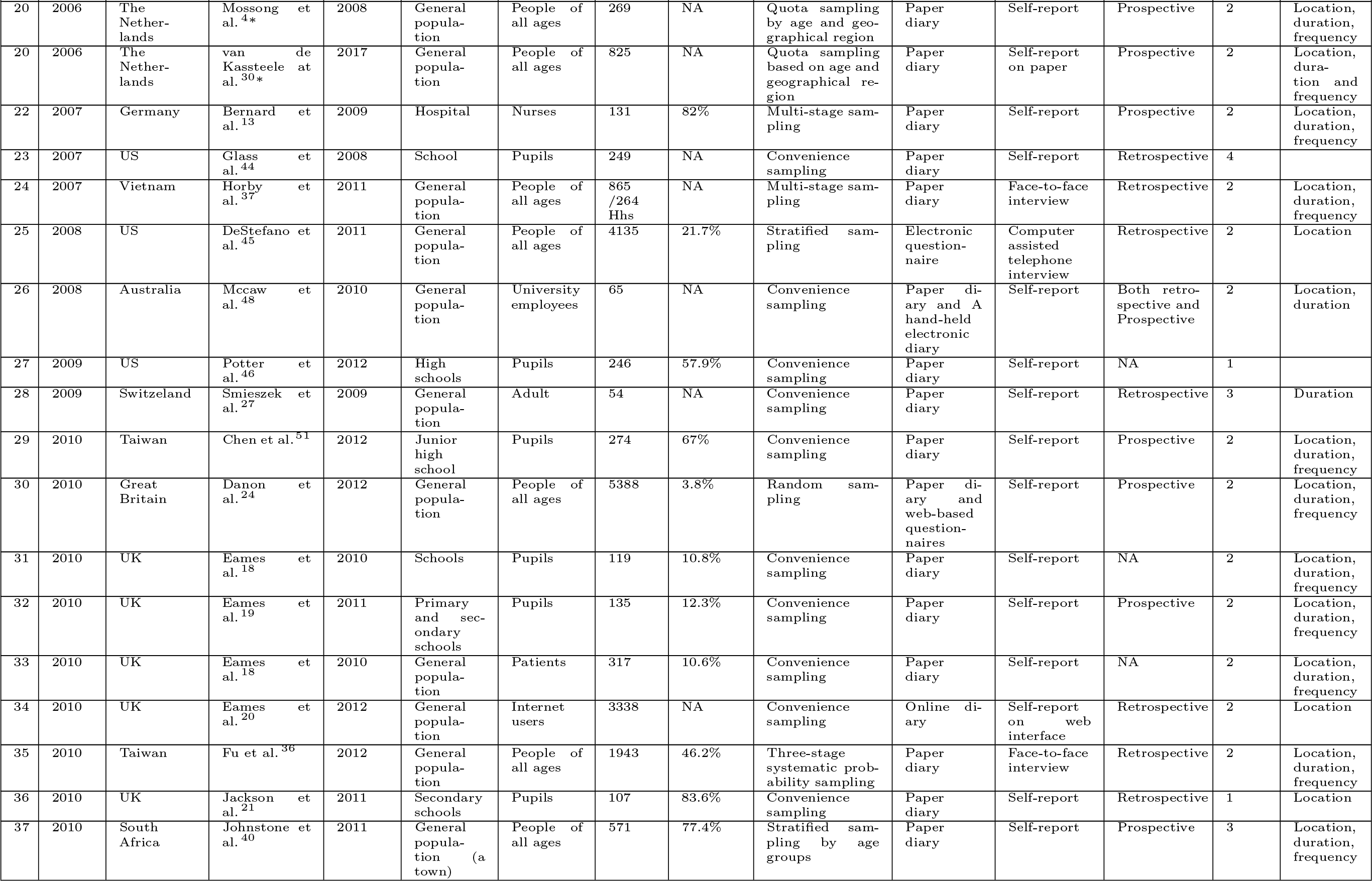
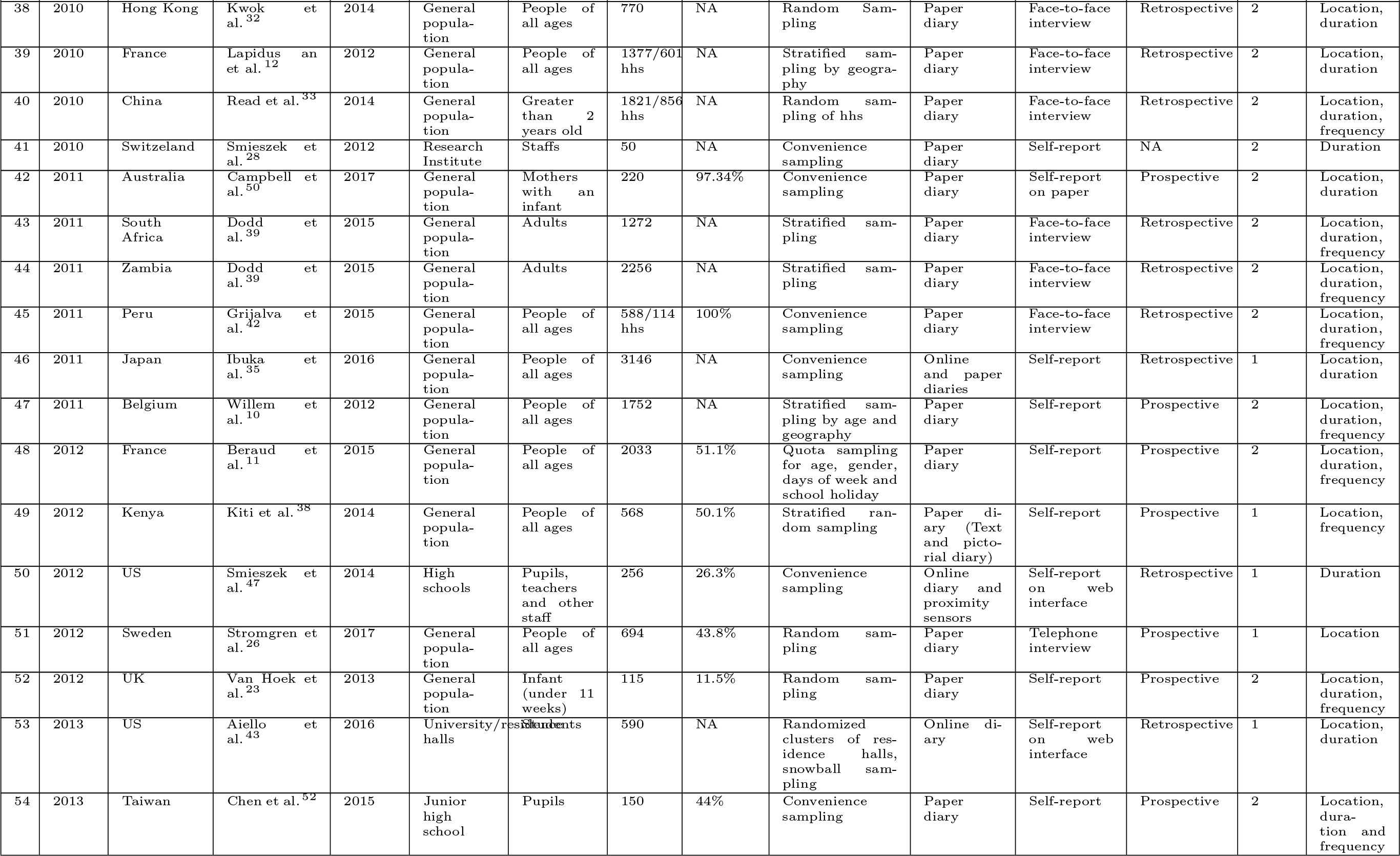
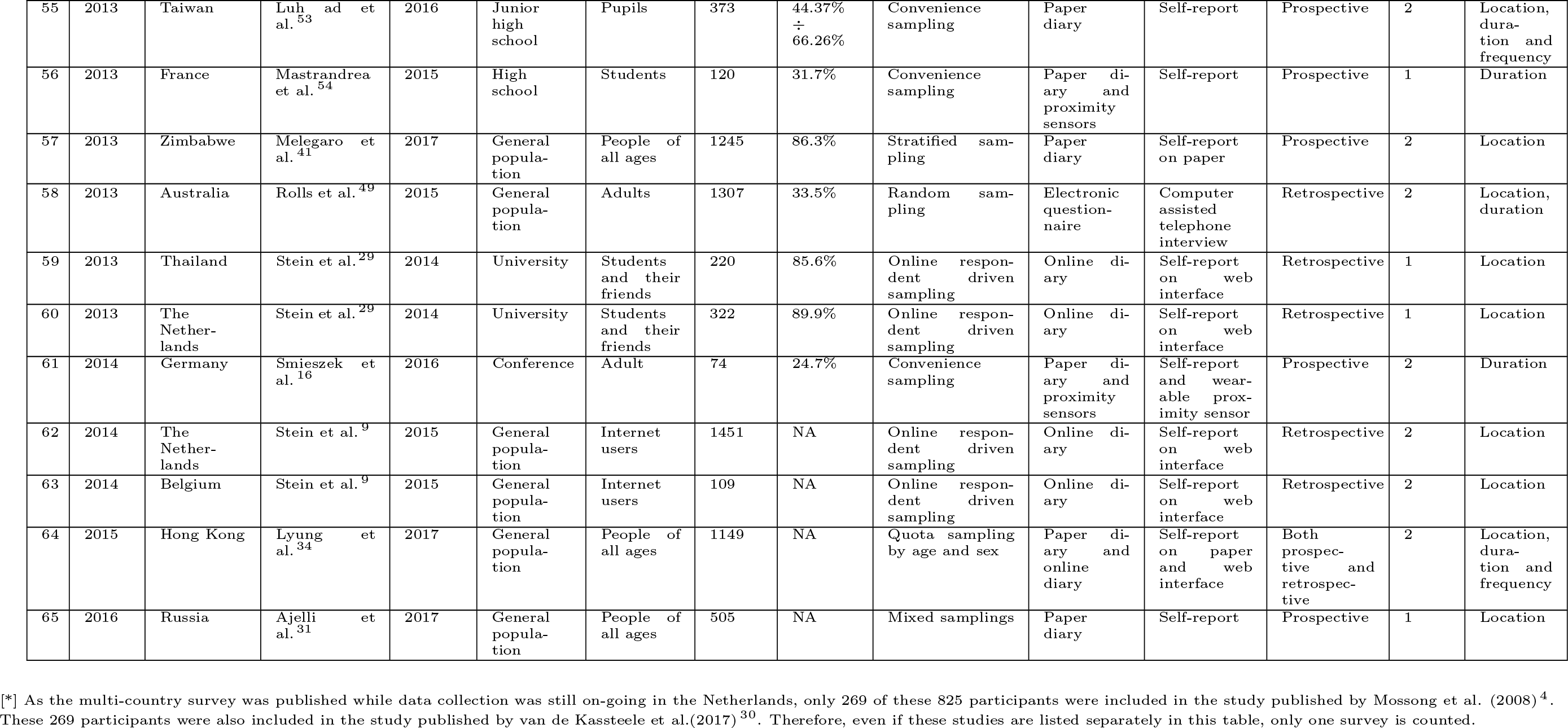
Summaries of social contact surveys.

## Discussion and conclusion

Social contact surveys are increasingly used to collect empirical data on human contact behavior and provide crucial inputs for mathematical models of infectious disease transmission. The POLYMOD project^4^ presented the first large scale representative population surveys conducted in 8 European countries. It also shared know-how both for data collection and analysis.

To date, most of these contact surveys were conducted in high and middle-income countries, whereas low-income countries, which have a higher burden of communicable diseases, were less studied in this respect. In view of this notion, there is a need to continue studying contact patterns more widely and in particular in low- and middle-income settings. It is also worth noting that in low- and middle-income countries, the choice to perform a general population representative survey may be less meaningful, given the large variety of different settings (urban, rural, etc) that are simultaneously present. Therefore, the decision for sampling specific areas can be deliberate and aimed at sampling social contact interactions in an area with distinct and more specific socio-demographics characteristics.

The use of an appropriate sample size is of importance to any study^56^. Most surveys did not clearly present sample size calculations, so we do not know to which extent important parameters, e.g., population size, confidence level and margin of error, were taken into account^61^. Sample size estimation is even more important when one wants to compare social contact surveys between or among populations. Given the lack of a clear picture regarding which demographical and anthropological factors are relevant in shaping contact patterns, inherent factors that may drive contact patterns are difficult to account for, thus making comparison among large populations, e.g. countries, even more difficult. In addition, the response rate or the substitution scheme also need to be considered to ensure that the number of study participants is obtained as initially planned. Indeed, the proportion of people who chose not to participate in surveys was quite high (on average 42.9%), particularly in general population-based surveys^11,20,23,24,45^. The extensive analysis of this review has underlined a general lack of information on response rates, and a call for a better non-response analysis emerges as a guideline for future studies.

The prospective design is subject to less recall bias than the retrospective design. This notion can be partly explained by the fact that respondents in the former are informed in advance about which days they will be assigned for reporting their contact information. Furthermore, they are also asked to keep a diary with them and finish reporting before the surveying day is elapsed. Thus, the prospective design requires more commitment from respondents. However, in return, a prospective design can obtain more reported contacts compared with retrospective design^15,48^. However, large-scale studies are needed to confirm this conclusion.

Each data collection tool has its own advantages and disadvantages^22^. For example, the use of paper or online contact diary is inaccurate for recording contacts of short duration, which are often forgotten by respondents^16,47,54,57^. On the other hand, proximity sensor devices can address this problem quite well. However, these devices only record contacts among people equipped with such devices; thus, it seems impractical to employ these devices in a large study population. The use of PDA seems to place more burden on study participants compared with a paper diary given that it necessitates them to input information about their contacts in an electronic device instead of just jotting down contacts in a paper diary^48^. In addition, this device may be difficult for those who are not familiar with the use of electronic devices. To date, comparison of the social mixing behavior captured by these data collection tools was only conducted in convenience surveys with small sample sizes^16,47,48,54,57^.

Self-reporting on paper or online diaries is the most commonly employed method in social contact surveys. Contact data collected by the self-report method were more complete compared with contact data collected by CATI because the former offers respondents more time to remember all contacts they made during a study period^58^. The use of the CATI method is primarily motivated by the cost-efficiency of the survey, allowing recruitment of a large and geographically well-defined population sample. However, one main disadvantage of a telephone interview lies in the fact that the duration of the telephone interview is probably too long given the complexity associated with verbal recall and repetitiveness for a diary type questionnaire, resulting in incomplete or missing information^49^. Perhaps, this is a main challenge that resulted in only two surveys using this method. Face-to-face interviews can help reduce inconsistencies of information provided by respondents in the presence of trained interviewers. In addition, this method also had a good response rate compared with other methods; however, the fieldwork and data collection are costly and time consuming^36,37,42^. To date, one study has compared contact behaviors collected by the self-report mode and face-to-face interview^3^.

The definition of a potentially infectious contact is of crucial importance given that it will be used as a surrogate for exposure to disease and help estimate transmission parameters^13^. Brankston et al.^62^ listed 4 modes of transmission of respiratory infections: direct contact, droplet, airborne and fomite or indirect contact. Direct contact and droplet require susceptible individuals to have close contact with infected individuals to enable infection, whereas airborne and fomite transmission do not^28^. Consequently, contact definitions in most surveys did not capture potential risks from the latter transmission modes^28^. Even for droplet transmission, the use of a face-to-face conversation definition to record non-physical contacts might lead to underreporting potentially infectious events given that susceptible individuals are likely able to contract a respiratory infection by just standing or sitting next to infected individuals who are sneezing, coughing, talking, etc. with no need to exchange words. Furthermore, it seems even more challenging to record common touching frequency of shared material objects, such as door knobs, water taps, public transport support poles, etc., for which two people do not need to be in each other’s physical proximity to be able to infect one another indirectly. Indeed, the more details on potentially infectious events we attempt to collect, the greater the burden we impose on respondents.

It is tempting to ask study participants to report their contacts as long as possible to obtain a broader view of contact patterns and gain insights in day-to-day variation. Nevertheless, the demanding and tedious task of diary keeping may prevent many participants from recording the information for a long time in prospective studies^63^. Beraud et al.^11^ demonstrated that participants reported 6% [1%-10%] less contacts on the second day of the survey. In addition, the more contacts they reported on the first day, the larger the proportional decrease in contacts on the second day. This finding might explain why only a few surveys asked respondents to report their contacts for a duration of more than 3 days^3,15,22,27,28,43^. For retrospective studies, a longer reporting period implies a longer recall period, with an associated larger bias. Therefore, in retrospective studies, researchers should try not to overstretch the reporting period.

The review provides information on the most relevant determinants of social contacts identified in previous studies. When designing future surveys, it is important to consider which characteristics may be sufficiently relevant to include as determinants. Asking study participants to report too many characteristics of contactees imposes a burden on participants. For example, collecting age of the participants and their contacts is informative, as some studies revealed that using age-related mixing patterns helped explain observed serological and infection patterns of infectious diseases like pertussis, varicella and parvovirus-B19^3,64,65^. In addition, collecting information about location, duration and frequency of contacts is also very essential for exploring mixing patterns of individuals and helping form effective strategies for disease prevention and control. Also, for school-aged children, a dominant number of contacts are made in school, leading to an indication that school closure can have a substantial impact on the spread of a respiratory infection^18,21,66–68^. When considering the duration and frequency of contact, it seems reasonable that a short random encounter of two persons is less likely to transmit a certain infectious disease compared with a more frequent encounter that lasts several hours. Accordingly, if all contacts are treated equally, it may lead to an overestimation of the individual transmission probability in cases of short, less frequent contacts and an underestimation in cases of long, more frequent contacts^27^. Several studies found that close contacts with a duration of at least 15 minutes involving skin-to-skin touching were most predictive of the pre-vaccination prevalence of Varicella Zoster Virus^64,69^. Therefore, age of the contactee and duration emerge as the most important information and should always be recorded in social contact surveys.

A comparison among all surveys based on a quantity such as the average number of contacts can be problematic. The sample serves as the first obstacle. Given different research questions or participant availability, not all the samples studied can be considered as representative of the envisaged population. This notion is important especially because age is a relevant determinant of social contacts, and samples in which a specific age class is over-represented regardless of study design can induce a strong bias on the number of contacts measured. For example, the study reporting the largest number of contacts (70.3^21^) was performed in a secondary school in the UK with the aim of estimating the reduction in social contacts due to school closure. Once these caveats are taken into account, Table 1 can be valuable to help identify all of the surveys sharing similar features that are relevant in addressing a specific research question. For example, the POLYMOD survey^4^ demonstrated that the main structure of social interaction among age categories was the same among several EU countries although the strength of the interaction could vary between countries. On the other hand, the average number of contacts measured in sub-Saharan countries by Dodd^39^ is considerably reduced compared with the average for high-income countries. In fact the development level can be important in determining social interactions, for example, due to local population density or reduced school attendance^41,42,69^. Quantifying the impact of different demographical factors on social contacts would require re-analysis of the datasets on the same basis and goes beyond the scope of this review. However, this re-analysis could be performed in the future as datasets of social contact surveys will be made available from researchers in a unified format^70^.

This review used PubMed and Web of Knowledge for searching publications, possibly resulting in the omission of relevant publications that can be captured from other online databases, e.g. Scopus or Embase. However, the availability and accessibility of PubMed and Web of Knowledge are of great value given that anyone can easily search for and refer to relevant articles that were used and cited in this review paper. In addition, the literature research step allowed us to recover more articles independently of a specific database, possibly recovering the ones we lost querying only PubMed and Web of Knowledge. We were exclusively interested in social contact surveys using the contact diary method, leading to the exclusion of a large number of surveys that exclusively employed direct observation or proximity sensor methods to measure contact behavior of individuals. The use of these methods along with their advantages and disadvantages have been summarized elsewhere^5,63^. Third, to the best of our knowledge, our search query omitted the relevant articles of Leecaster et al.^57^ and Kwok et al.^71^, which are eligible for this review. These articles are missing the survey/questionnaire/diary word in the abstract and title and therefore elude our searching method. The recent publication date (2016 and 2018) also prevented these articles from appearing in the references of the relevant articles.

Since the POLYMOD survey, there has been an increasing trend in the number of social contact surveys used to collect empirical contact data. Social contact surveys have been conducted widely in many countries, but most focused on high-income countries. These surveys used a range of different study designs with different study subjects, settings, sampling scheme to study designs, data collection tools and data collection methods. Moreover, the definition of “contact” and its characteristics also differ, making comparison of contact patterns among surveys even more difficult. Improvements towards a unified definition of “contact” and standard practice in data collection could help increase the quality of collected data, leading to more robust and reliable conclusions about contact patterns of individuals.

This review demonstrates that contact surveys typically have in the order of a thousand participants, rely on convenience sampling, and use a retrospective design with paper diaries and self-reporting of contacts over a single day. Major determinants for this number of contactees include characteristics of the respondent (age, gender, and health status), time (weekday or weekend, and term time or holiday) and their immediate environment (household size and urban versus rural). A typical number of different contactees reported per day is in the order of 20 for country wide studies, a quantity that proved remarkably robust despite the many different study designs.

## Supporting information

**S1 File. The comprehensive table of all social contact surveys**.

**S1 Fig. The average samples**. Average sample size of surveys in general population and surveys in specific setting.

**S2 Fig. Sampling schemes by continents and periods**.

**S3 Fig. Sampling schemes in details**.

**S4 Fig. Distribution of social contact surveys by study designs**.

**S5 Fig. Average number of contacts by factors**. Average number of contacts measured and 95% CI in weekday vs weekend (a), holiday vs. term-time (b), well vs unwell health status (c) and during vs outside flu season (d). For surveys reporting mean number and standard deviation a 95% CI for the mean has been computed.

**S6 Fig. Median of number of contacts among surveys**. Median and quantiles of contacts. Surveys are labeled according to the publication’s first author, year and the country in which the survey was performed.

**S7 Fig. Global map of social contact surveys**.

## Acknowledgments

This work is part of a project that has received funding from the European Research Council (ERC) under the European Unions Horizon 2020 research and innovation programme (grant agreement 682540 TransMID). AM acknowledges the ERC project DECIDE (ERCStG283955).

## Author Contributions

TVH, PC, PB and NH conceived and designed the inclusion criteria. TVH and PC performed the screening. TVH, PC and NH wrote a first draft of the paper. TVH, PC, AM, JW, CG, PB and NH contributed to the final version of the paper. All authors read and approved the final manuscript.

